# Exploring a novel genomic safe-haven site in the human pathogenic mould *Aspergillus fumigatus*

**DOI:** 10.1101/2022.03.28.486018

**Authors:** Takanori Furukawa, Norman van Rhijn, Harry Chown, Johanna Rhodes, Narjes Alfuraji, Rachael Fortune-Grant, Elaine Bignell, Matthew C. Fisher, Michael Bromley

**Author notes:** These authors contributed equally (Takanori Furukawa, Norman van Rhijn). **To whom correspondence should be addressed:** Michael J Bromley.

## Abstract

*Aspergillus fumigatus* is the most important airborne fungal pathogen and allergen of humans causing high morbidity and mortality worldwide. The factors that govern pathogenicity of this organism are multi-factorial and are poorly understood. Molecular tools to dissect the mechanisms of pathogenicity in *A. fumigatus* have improved significantly over the last 20 years however many procedures have not been standardised for *A. fumigatus*. Here, we present a new genomic safe-haven locus at the site of an inactivated transposon, named SH-*aft4*, which can be used to insert DNA sequences in the genome of this fungus without impacting its phenotype. We show that we are able to effectively express a transgene construct from the SH-*aft4* and that natural regulation of promoter function is conserved at this site. Furthermore, the SH-*aft4* locus is highly conserved in the genome of a wide range of clinical and environmental isolates including the isolates commonly used by many laboratories CEA10, Af293 and ATCC46645, allowing a wide range of isolates to be manipulated. Our results show that the *aft4* locus can serve as a site for integration of a wide range of genetic constructs to aid functional genomics studies of this important human fungal pathogen.

## INTRODUCTION

*A. fumigatus* is a saphrotrophic filamentous fungus that can cause a spectrum of clinical diseases ranging from allergic sensitisation and chronic infection to life-threatening invasive infection depending on the host immune status (Kosmidis and Denning 2015). Worldwide, infections caused by this cosmopolitan fungus lead to in excess of 600,000 deaths every year (Bongomin et al. 2017) and our ability to treat these infections remains critically challenged as there are limited treatment options. The development of robustly validated methodologies to fully explore genetically encoded mechanisms that contribute to virulence of *A. fumigatus* is central to our efforts in developing novel treatments.

Over the last 20 years, a variety of molecular tools have been developed to genetically manipulate *A. fumigatus* (Krappmann 2006; Weld et al. 2006; Morio et al. 2020; Van Rhijn et al. 2020, Zhao et al. 2019) and we are now readily able to replace genes, promoters, introduce fluorescent and immunogenic tags and create point mutations in laboratory strains and environmental isolates. Confirmation that any changes implemented in the genome are solely attributed to the intended mutation is usually achieved by reverting the genotype by performing a second genetic manipulation. However, there are currently no standardized approaches for this type of complementation. Often targeted integration of a large complementation cassette back to the mutated locus is performed. Unfortunately, the need to co-integrate a selectable marker under the control of a highly expressing and constitutive promoter may result in the dysregulation of the target gene, its neighboring genes or proximal non-coding RNAs resulting in unforeseen phenotypic changes (Verdoes et al. 1995; Fleißner and Dersch 2010).

One method to overcome this is to identify a defined genetic locus for integration of endogenous and exogenous DNA. The concept of “genomic safe-harbor sites” (GSHs) is originally derived from higher eukaryotes and is defined as a region of the genome that is considered to be both transcriptionally active and its disruption does not lead to any discernible phenotypic effects (Zambrowicz et al. 1997; DeKelver et al. 2010; Sadelain et al. 2012; Li et al. 2014; Papapetrou and Schambach 2016). GSHs are ideal integration sites for basic functional genomics research as well as for gene therapy applications. Similar genetic loci, which are known as “genomic safe-haven sites” (SH), have recently been identified and used successfully in a number of human pathogenic fungi (Arras et al. 2015; Upadhya et al. 2017; Fan and Lin 2018; Fan and Lin 2020). Recently, two potential genomic safe-haven regions SH1 and SH2 were identified in *A. fumigatus* based on genomic annotation and transcriptome data profiling (Pham et al. 2020). The authors found that insertion of DNA at the SH1 locus did not cause obvious changes in the expression of neighboring genes or impact growth. Furthermore, active expression of a heterologous fluorescence protein was demonstrated from these two safe-haven regions. Although data was presented showing this locus did not impact virulence in a wax moth (*Galleria mellonella*) model, no information was presented about virulence in mice.

In this study, we have identified a new genomic safe-haven, that we term SH-*aft4,* at the site of an inactive Tc1/mariner type transposable element (*aft4*). Our analyses revealed that insertion of a gene expression cassette at SH-*aft4* does not have any significant impact on growth, cytotoxicity, or pathogenicity of *A. fumigatus* in either *G. mellonella* or murine pulmonary infection models. Furthermore, we show that SH-*aft4* is highly conserved in the genomes of many clinical and environmental isolates of *A. fumigatus*. We hence provide evidence that SH-*aft4* locus can be used as a safe-haven site for gene complementation and transgene expression to aid functional genomics studies in *A. fumigatus*.

## MATERIALS AND METHODS

### Ethics statement (for PLOS Journals)

The murine infection experiments were performed under UK Home Office Project Licence PDF8402B7 and approved by the University of Manchester Ethics Committee.

### Strains, media and culture conditions

*A. fumigatus* MFIG001, a member of the CEA10 lineage lacking a functional *ku80* gene (Fraczek et al. 2013; Bertuzzi et al. 2020), was used as the parental strain for transformation throughout this study. Strains were cultivated on Sabouraud dextrose agar (SAB) and conidia were harvested in 0.1% Tween-20. For growth phenotyping on solid medium, 1×10^3^ of conidia were spotted on Aspergillus Minimal Medium (AMM) or Aspergillus Complete Media (ACM) (Pontecorvo et al. 1953) and incubated at 37 °C for 72 h. For Western blotting and quantitative reverse transcription PCR (qRT-PCR) analysis, fungal strains were cultivated at 37°C for 24 h in AMM with 1% (w/v) glucose as a carbon source and 20 mM ammonium tartrate as a nitrogen source. Iron was omitted to mimic iron starvation (-Fe). For iron-replete conditions, 30 μM FeSO_4_ was added to AMM (+Fe). Mycelia were collected by filtration, immediately frozen with liquid nitrogen, and kept at -80°C until use.

### PCR amplification of the *aft4-*encoding genetic locus in the commonly used *A. fumigatus* laboratory strains

Total genomic DNA was extracted from the spores of the *A. fumigatus* strains (Af293, CEA10, ATCC46645, and AfS35) using a standard CTAB DNA extraction method (van Burik et al. 1998). The presence of the *aft4* locus in the respective *A. fumigatus* strains was investigated by amplifying different regions of the *aft4* transposon by PCR. Primers used for this analysis were designed from the genome sequence of *A. fumigatus* A1163, and are listed in Supplementary Table 1.

### Generation of *aft4-hyg* knockout mutant

The gene replacement cassette for the *aft4* locus was generated using fusion PCR and transformed into *A. fumigatus* MFIG001 as described previously (Zhao et al. 2019). Briefly, about 1 kb of the 5’-upstream and the 3’-downstream regions of the *aft4* transposon unit were amplified by PCR and fused with the hygromycin B phosphotransferase (*hph*) resistance marker cassette using the primers aft4 P1 to P6 and hph Fw and Rv (Supplementary Table 1). Single copy integration of the gene replacement cassette at the *aft4* locus was verified by Southern blotting analysis with PvuI and BstXI using 0.7 kb of the *hph* fragment as a probe (Supplementary Fig. 3).

### Germination rate and hyphal extension analysis

Germination rate and hyphal extension rate were determined as follows. 500 µl of 5×10^5^ spores of the wild-type (MFIG001) or the *aft*4-hyg mutant was inoculated in RPMI-1640 medium containing 2.0 % glucose and 165 mM MOPS buffer (pH7.0) in a 24 well glass bottom plate. The culture was incubated at 37 °C and images were taken on a Leica SP8X confocal microscope (Leica). Germination rate and hyphal length were measured in ImageJ by scoring images manually or measuring hyphal length by using the segmented tool, respectively.

### Construction of the NLS-venus expressing mutants

The plasmid phapXNLS-venus (Supplementary Fig. 4) and the NLS-Venus expressing strain (Supplementary Fig. 3b, NLS-venus cassette1) was constructed previously (Furukawa et al. 2020) using *A. fumigatus* MFIG001 as a parental strain (van Rhijn et al. 2020). In the NLS-Venus expressing strains, a yellow fluorescent protein derivative Venus (Nagai et al. 2002) encoding gene was fused with a nuclear localization signal (NLS; PKKKRKV) derived from the Simian Virus 40 (SV40) large T-antigen. The construct was placed under control of the *hapX* promoter.

The NLS-venus expression cassette with the hygromycin resistance marker (Supplementary Fig. 4b, NLS-venus cassette2) was amplified by PCR from phapXNLS-venus with the primers Hyg-hapXP-NLS-Venus P7 and hapXP-NLS-Venus P6. The resultant NLS-venus expression cassette contains 50-bp of homology arms to facilitate homologous recombination, and the expression cassette was introduced to the *atf4* locus of *A. fumigatus* MFIG001 by using a CRISPR-Cas9 mediated genome-editing system (Al Abdallah et al. 2017; van Rhijn et al. 2020). Correct integration of the expression cassette was verified by PCR and Sanger sequencing. The primers and guide RNAs used for the generation of the NLS-venus expressing strains are listed in Supplementary Table 1. The details of the genome-editing at the *aft4* locus are also shown in Supplementary Figure 6.

### Western blotting analysis

Mycelia of the NLS-venus expressing strains were frozen in liquid nitrogen and ground to a fine powder before incubation on ice in 1.0 mL of ice-cold cell-lysis buffer (50 mM HEPES-KOH pH 7.5, 150 mM NaCl, 1 mM EDTA, 1% Triton X-100, 0.1% deoxycholate (Sigma D6750), 0.1% SDS, 1 mM PMSF and fungal proteinase inhibitor cocktail (Sigma)) for 10 min. Mycelia were lysed with a FastPrep bead beater (MP Biomedicals) for 1 min at 6,000 rpm, and the lysate was further incubated on-ice for 10 min. Soluble protein fraction was recovered by centrifugation and then precipitated using 10 % trichloroacetic acid (TCA). 100 µg of the TCA precipitated protein were resolved on an SDS-PAGE gel, transferred to a PVDF membrane, and subjected to western blotting. A polyclonal anti-GFP-antibody (A-11122, Thermo Fischer Scientific) was used with a horseradish peroxidase-conjugated secondary anti-rabbit IgG (Ab6721, Abcam) and Clarity Western ECL substrate (Bio-rad) to detect the expression of NLS-Venus protein. Images were taken by exposing the blots to the ChemiDoc XRS gel imaging system (Bio-rad)

### Fluorescent microscopy

1×10^2^ spores of the NLS-Venus strains were grown for 24 h at 37 °C in 200 μL of AMM with different iron conditions in an 8 well chamber (Ibidi). Fluorescent and bright field live-cell imaging were performed using a Leica TCS SP8 confocal laser scanning microscope equipped with hybrid GaAsP (HyD) detectors and a 60 × objective. Argon laser 488 nm was used for fluorescence excitation. Confocal microscopy images were analysed and processed with ImageJ (Schneider et al. 2012).

### Quantitative reverse transcriptase PCR

Total RNA was extracted using TRI reagent® (Sigma-Aldrich) according to the manufacturer’s instructions. The extracted RNA samples were treated with RQ1 RNase-Free DNase (Promega) and further purified using the RNeasy Mini kit (Qiagen). First-strand cDNA was synthesized using the iScript cDNA synthesis kit (Bio-Rad) with 100 ng of total RNA as a template. Amplification reactions were performed using the iTaq Universal SYBR Green Supermix (Bio-Rad) in a final volume of 20 µL using 0.5 µM forward primer, 0.5 µM reverse primer, and 2 µL of 5-fold diluted cDNA as a template. Primers used in this analysis are listed in Supplementary Table 1. The relative expression level of the transcripts of the neighboring genes was determined using the ΔΔCt method with the housekeeping glyceraldehyde-3-phosphate dehydrogenase encoding *gpdA* gene, as a reference. The specificity of the PCR reactions was documented by melting curve analysis. Experiments were performed in biological triplicates. A Mann Whitney U test was used for statistical analysis (p <0.05 considered statistically significant).

### Cell toxicity assays

A549 human pulmonary carcinoma epithelial cells (American type culture collection, CCL-185) were used for cell toxicity assay under passage 20. Cells were maintained at 37 °C, 5% CO_2_ in Dulbecco’s Modified Eagle’s Medium (DMEM), 10 % foetal bovine serum (FBS), 1 % Penicillin-Streptomycin (Sigma-Aldrich). For all experiments, 2×10^5^ A549 cells were seeded in 24-well plates and incubated for 16 h when confluence equals 90 %. Cells were then challenged with 10^5^ spores of the wild-type strain or the *aft4-hyg* knockout mutant and incubated for 24 h. Following co-incubation with *A. fumigatus* spores, cell culture supernatants were collected, and the level of cell toxicity was measured using a lactate dehydrogenase assay kit (Promega). Experiments were performed in five biological replicates and technical duplicate (n=10). Uninfected cells were used as a negative control. Statistical analysis was carried out using GraphPad Prism 8.0 (La Jolla, CA, USA).

### Galleria mellonella infection model

The *G. mellonella* infection model was performed as described in Johns et al 2017. Briefly larval *G. mellonella* (Live Foods Company (Sheffield, England)) (Minimum weight 0.3g; 10 per group) were inoculated by injecting 10 µl of a 1 × 10^6^ spores/ml suspension into the last, left proleg. Sham infections were performed with PBS. Larvae were monitored daily for 8 days and scored for mortality.

### Murine infection models

The murine infection experiments were performed under UK Home Office Project Licence PDF8402B7 and approved by the University of Manchester Ethics Committee. *A. fumigatus* strains were cultured on ACM containing 10 mM ammonium tartrate for 6 days at 37 °C and conidia were harvested in sterile saline, washed twice in sterile saline, and used for infection experiments.

CD1 male mice (Charles <u>River UK, L</u>td.) (19-25g) were housed in groups of 3-4 in IVC cages with access to food and water *ab libitum*. All mice were given 2 g/L neomycin sulphate in their drinking water throughout the course of the study. For the pulmonary model of infection, mice were rendered leukopenic by the administration of cyclophosphamide (150 mg/kg of body weight; intraperitoneal) on days -3, -1, +2, and every subsequent third day, and a single subcutaneous dose of cortisone acetate (250 mg/kg) was administrated on day -1. Mice were anaesthetized by exposure to 2-3 % inhalational isoflurane and infected intranasally with a spore suspension of 1.25 x 10^7^ conidia in 40 µl of saline solution. Mice were weighed every 24 h from day -3, relative to the day of infection, and visual inspections were made twice daily. In the majority of cases, the endpoint for survival in experimentation was a 20% reduction in body weight measured from day of infection, at which point the mice were sacrificed. Kaplan-Meier survival analysis was used to create a population survival curve and to estimate survival over time, and *p*-values were calculated through a log rank analysis.

### Bioinformatic analysis of the *aft4* locus in the genome of various *A. fumigatus* isolates

A total of 218 assembled and annotated genomes alongside a pan-genome and phylogenetic data from a previous study were acquired for the identification of *SH-aft4* homologues (Sewell et al. 2019; Rhodes et al. 2021). Terminal inverted repeat (TIR) regions were first aligned to the genomes, followed by the coding sequence of *aft4,* using BLAST, v2.0.9, (E-value < 0.01). Next, we identified TIRs within 10kb of *aft4*, and extracted all nucleotide sequences, using BEDTools, v2.30.0, intersect and closest. The same software was then used to identify neighbouring genes of each TIR*-aft4-*TIR (*SH-aft4*) homolog and cross-referenced with the pan-genomic data to identify conserved syntenic regions. The *aft4* region from each *SH-aft4* locus underwent global alignment to the putative *aft4* from Af293 using EMBOSS, v6.6.0.0, Needle. Furthermore, the largest translated ORF was identified within each of these regions using “getorf” (EMBOSS), samtools, v1.3.1, and seqtk, v1.3-r106, and also aligned to Aft4, using Needle. Percentage identities from the pairwise alignments were then extracted. Multiple sequence alignments were produced for *NH1-aft4 and NH2-aft4,* and their protein products, using MUSCLE, v3.8.1551. Sequence alignments were then annotated and view in Jalview, v2.11.1.7. Previously generated phylogenetic data, including clade structure, was combined with *SH-aft4* loci presence and visualised using iTOL, v6.5.2.

## RESULTS

### Identification of the *aft4* locus, encoding an inactive transposable element, as a potential genomic safe-haven site in *A. fumigatus*

We have previously explored the genome of *A. fumigatus* Af293 to identify potential functional transposable elements in an attempt to develop molecular mutagenesis tools for this fungus (Hey 2007; Hey et al. 2008). During the genomic survey, a unique 1.3 kb nucleotide region (Chr6: 2,227,032-2,228,286) was identified as an element with a moderate homology (66%) to the *Fusarium oxysporum* impala (Langin et al. 1995), a member of the *Tc1/mariner* superfamily of class II transposons (Daboussi and Capy 2003). Analysis of this locus revealed that the element, we name here as *aft*4 (*<u>A</u>. fumigatus* transposon <u>4</u>), possesses a small putative open reading frame (ORF) with 158 amino acids that has a similarity (55%) to the C-terminal part of the *F. oxysporum* impala transposase, and is flanked by 29 bp perfect terminal inverted repeats (TIR) (Supplementary Fig. 1a-c). A complete ORF of 348 aa, which is likely to represent an entire functional transposase appears to have been interrupted by point mutations (Fig.1a and Supplementary Fig. 1bc) which introduces two premature stop codons and disrupts the catalytic centre consisting of the DDE triad (Supplementary Fig1c). We hypothesized that if the *aft4* ORF is no longer functional, it would be possible to use this genetic locus as a genomic safe-haven site in *A. fumigatus*.

**Figure 1.**
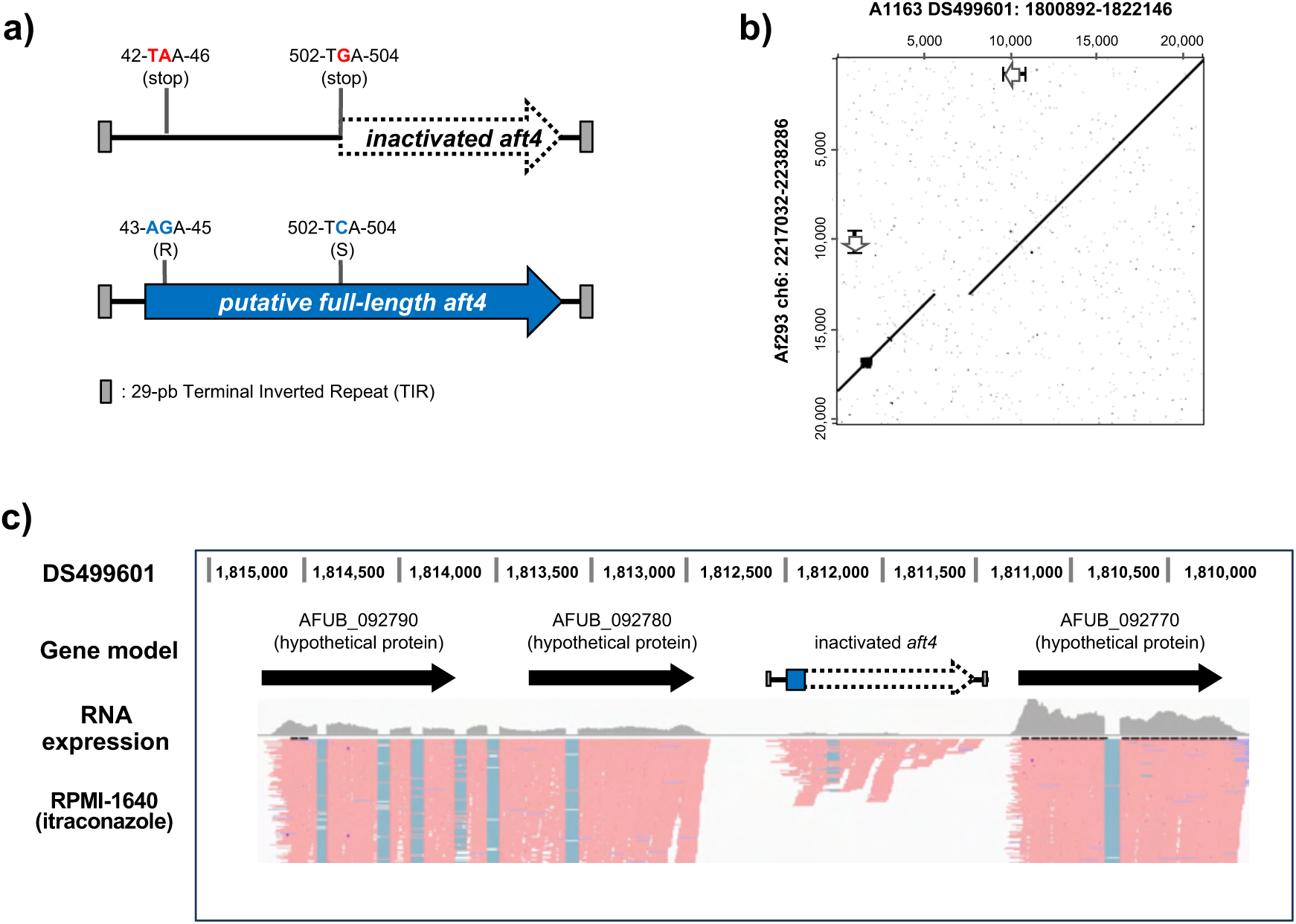
Identification of the *aft4* locus as a potential safe-haven site in *A. fumigatus*. **(a)** Schematic representation of *aft4* encoding a putative inactivated Tc1/mariner like transposable element. The dotted arrow represents the putative inactivated form of the *aft4* ORF found in the *A. fumigatus* genome, and the blue arrow represents the putative full-length from of the *aft4* ORF. The full-length ORF can be generated from the inactivated *aft4* by introducing two nucleotide changes (T(43)A and G(503)C) as shown in the figure. The gray boxes represent the terminal inverted repeats (TIRs). **(b)** Dot plot analysis of the 20-kbp region surrounding the *aft4* element between *A. fumigatus* Af293 and A1163. The white arrows represent the genomic coordinate of the inactivated *aft4* ORF. **(c)** RNA-expression profile of the *aft4* locus in *A. fumigatus* A1163. The strain was pre-cultured in AMM for 37°C for 18h, and then transferred into RPMI-1640 + 0.5 mg/L itraconazole for 4 h. Reads were visualized using IGV.

We investigated the presence of the *aft4* transposon unit in the publicly available genome sequence of the *A. fumigatus* strains, Af293 and A1163, at FungiDB (https://fungidb.org/fungidb (Stajich et al. 2012)). A BLASTN search using the *aft4* sequence derived from *A. fumigatus* Af293 as a query showed that *aft4* is present as a single copy element in the genome of A1163 with 99% identity. All the nucleotide changes that cause the nonsense mutations within the *aft4* ORF were conserved in both strains, suggesting that *aft4* in the A1163 genome also encodes a non-functional transposable element. The 20-kbp regions surrounding the *aft4* locus share a high degree of sequence identity between the two genomes (Fig. 1b) with the exception of a c. 2kb DNA fragment (DS499601:1806511:1808504) in genome of A1163 which is not present in the same region of the genome of Af293. This 2kb fragment appears to be a second transposon that has inserted 2.4kb bases downstream of *aft4* and can be found in at least 6 copies in the A1163 genome and 7 copies in the Af293 genome. These results demonstrate that *aft4* is syntenic in the Af293 and the A1163 genomes. As Af293 and CEA10, the parental isolate of A1163, belong to different genetic clades of *A. fumigatus* (Garcia-Rubio et al. 2018; Bertuzzi et al. 2020), we postulated that the *aft4* element is also present and fixed in the genome of the other *A. fumigatus* strains. To assess this possibility, we designed several pairs of primers to amplify different genomic regions surrounding *aft4* and examined the presence of the *aft4* element in the genome of the *A. fumigatus* isolates in common laboratory use (Bertuzzi et al. 2020) by PCR (Supplementary Fig. 2). Among the four tested *A. fumigatus* strains (*A. fumigatus* Af293, CEA10, ATCC46645, and D141), three of them showed positive amplification for the *aft4* element and its immediate flanking regions. Only *A. fumigatus* D141, which was originally isolated from an aspergilloma (Staib et al. 1980), did not show any amplification for all the tested pairs of primers.

Transposable elements and their surrounding regions are often targeted by fungal derived protective mechanisms that introduce epigenetic silencing (Daboussi and Capy 2003; Castanera et al. 2016). If this site were to be used as a safe-haven we would need to demonstrate that the locus is transcriptionally active. The genomic annotation derived from the *A. fumigatus* A1163 indicate that *aft4* is located in between two putative ORFs (AFUB_092770 and AFUB_092780), which are arranged in the tandem orientation (Fig. 1c,). Despite the fact that this locus does not seem to encode a functional transposon, analysis of our previous RNA-seq data (Gsaller et al 2016) revealed that the region is transcribed (Fig. 1c). Interesting the *aft4* transcript appears to include an intron. Even when this intron is spliced out however, no full ORF can be discerned. The two neighbouring genes were also found to be actively expressed, indicating that the *aft4* locus lies in a transcriptionally active genomic region.

### Deletion of the inactivated *aft4* element does not have significant impact on growth characteristics of *A. fumigatus*

Given the potential of the *aft4* locus as a genetic safe-haven site, it was important to assess whether the deletion of the inactivated *aft4* ORF or the insertion of a transgene expression cassette at the *aft4* locus has any impacts on growth of *A. fumigatus*. To this end a gene knockout mutant (*aft4-hyg*) was constructed in the *A. fumigatus* MFIG001 genetic background by replacing the *aft4* element with the hygromycin B resistance marker cassette (Fig. 2a). In this gene knockout cassette, expression of the hygromycin resistance gene (*hph* encoding a hygromycin B phosphotransferase from *Escherichia coli*), is under the control of the *A. nidulans gpd* promoter, which is frequently used for high level constitutive expression of transgenes in the *Aspergilli*. A strain carrying the *hph* marker was isolated by selection on hygromycin and single copy integration at the *aft4* site was confirmed in this strain by Southern blotting (Supplementary Fig. 3) highlighting the ability of the *aft4* locus to tolerate integration of a transgene.

**Figure 2.**
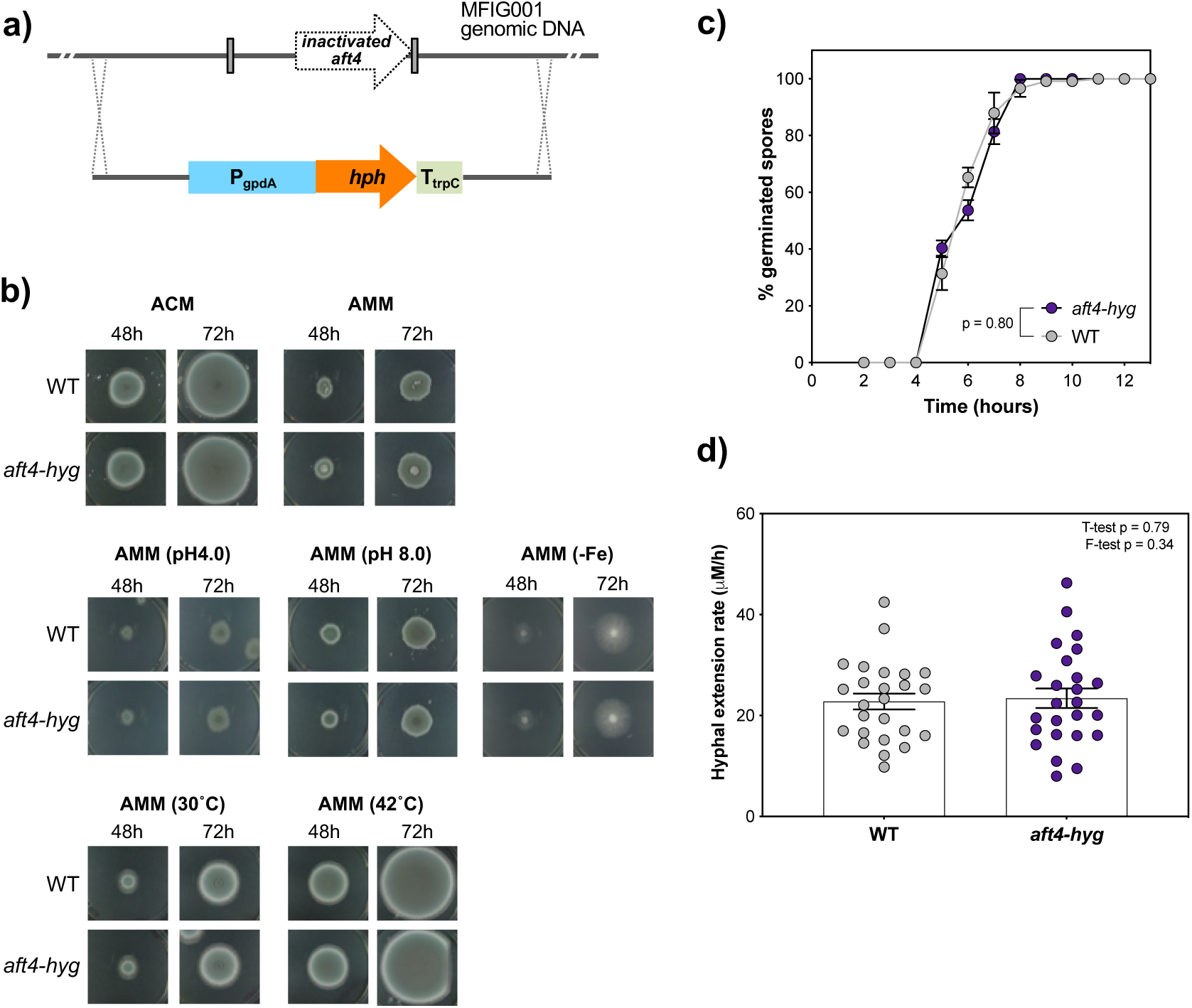
Deletion of the *aft4* element does not affect growth characteristics of *A. fumigatus*. **(a)** Schematic overview of the construction of the *aft4-hyg* knockout mutant. **(b)** Colonial growth phenotypes of the wild-type (MFIG001:WT), and the *aft4* knockout mutant *(aft4-hyg)* grown on a solid ACM or solid AMM with an infection relevant stress. **(c)** Germination profiles of the strains in a liquid RPMI-1640 medium. *p*-value was calculated by repeated measures 2way-ANOVA with Sidaks correction: NS, *P*>0.05. **(c)** Hyphal extension rates in a liquid in RPMI-1640 at 37 °C. *p*-value was calculated by Kruskal-Wallis test with Dunn’s correction: NS, *P*>0.05. Percentages of germination rates and hyphal extension rates were measured under the microscope.

We assessed the impact of the *aft4* deletion by characterizing growth phenotypes of the *aft4-hyg* replacement mutant alongside the parental strain MFIG001 (WT) on solid Aspergillus Complete Medium (ACM) and Aspergillus Minimal Medium (AMM). We also examined growth characteristics of *aft4-hyg* under several infection relevant stress conditions, which include acidic and alkaline pH, iron limitation, and high temperature (Seyedmousavi et al. 2015). No significant differences were observed in radial growth rates or colony morphology between *aft4-hyg* and MFIG001 in the tested culture conditions (Fig 2b).

Next, we examined the effects of *aft4* replacement on conidial germination and hyphal extension rate by culturing the mutant in a liquid RPMI-1640 medium (Fig. 2 c and d). After 5 h of incubation at 37 °C, conidia of the wild-type and *aft4-hyg* started to germinate and the germination rate reached almost 100 % after 8 h (Fig. 2c). The wild-type and *aft4-hyg* showed nearly identical germination patterns and no significant differences were detected between the strains (*p* = 0.80). When the hyphal extension rate was measured, a significant heterogeneity was observed within the population of the wild-type strain (Fig. 2d). This is consistent with the recent description of asynchronous germination of *A. fumigatus* spores and initial growth rates being an inherent property of each conidium (Danion et al. 2021; Earl Kang et al. 2021). Importantly, this heterogeneous hyphal extension pattern was conserved within the population of the *aft4-hyg* conidia, and no major differences were detected compared to the wild-type strain (T-test *p* = 0.79, F-test *p* = 0.34). Overall, these results indicate that deletion of the *aft4* element and expression of a transgene from the *aft4* locus have no significant impact on growth characteristics of *A. fumigatus*.

### Deletion of the inactivated *aft4* element does not have significant impact on the pathogenicity of *A. fumigatus*

To examine the impact of the gene replacement at the *aft4* locus on the virulence of *A. fumigatus*, we characterized the pathogenic traits of the *aft4-hyg* mutant using both *in vitro* and *in vivo* infection models. As spores interact with lung epithelial cells at the early stages of infection (Bertuzzi et al. 2018; Bertuzzi et al. 2019), we co-cultured the deletion mutants with human A549 lung epithelial cells. Cell damage was quantified by measuring the released lactose dehydrogenase (LDH) activity after 24 h of the infection challenge. As shown in Fig. 3a, the *aft4-hyg* strain showed similar levels of cytotoxicity compared to the parental strain and they were statistically indistinguishable from each other (*p* = 0.29). We examined the virulence of the *aft4-hyg* and isotype control strains in a *Galleria mellonella* infection model. No discernable differences were observed between the two strains. We next examined the virulence of the *aft4-hyg* strain in a leukopenic model of invasive aspergillosis. In this model, all mice challenged with the parental strain succumbed to infection (100% mortality) within 4 days post-challenge. Consistent with the observed growth characteristics (Fig. 2) and the unchanged level of cytotoxicity of the *aft4-hyg* strain (Fig. 3a), no significant differences were observed in the virulence of the *aft4-hyg* mutant compared to the wild-type in this murine infection model (*p* = 0.35) (Fig. 3b). Overall, our results indicate that deletion of *aft4* and the constitutive expression of the *hph* selection marker gene at this site does not have any significant impact on the pathogenic properties of *A. fumigatus*.

**Figure 3.**
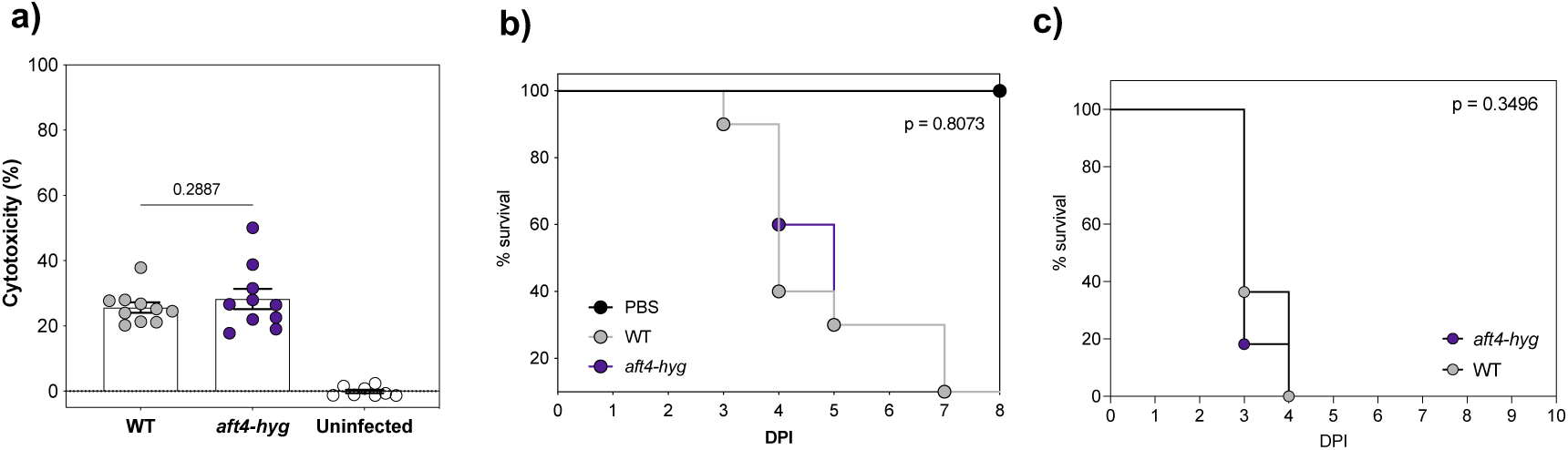
Deletion of the *aft4* element does not impact on the cytotoxicity and the pathogenicity of *A. fumigatus*. **(a)** Cytotoxicity of the wild-type (WT) and the *aft* knockout mutant (*aft4-hyg*). A549 epithelial cells were infected with the strains for 24 hours and their cytotoxicity was evaluated by measuring the release of lactose dehydrogenase (LDH) activity into the culture medium. The data represents 10 different infection challenges with triplicate LDH activity measurements. Cytotoxicity is expressed as a relative percentage of the released LDH activity after infection where complete lysis of cells is set equal to 100%. The error bars mean the standard error of the mean (SEM), and *p-*values were calculated by Kruskal-Wallis test with Dunn’s correction. **(b)** Effect of the deletion of the aft4 elements on virulence of *A. fumigatus* in a *Galleria mellonella* model of infection. A log rank analysis was used to compare the results between the strains. **(c)** Effect of the deletion of the *aft4* element on virulence of *A. fumigatus* in a murine model of invasive pulmonary aspergillosis. Kaplan–Meier curve for murine survival infected with the wild-type (WT), and the *aft4* knockout mutant (*aft4-hyg)* are shown. Mice were rendered neutropenic by treatment with cyclophosphamide and challenged with 5.0 × 10^5^ spores via intranasal route. A log rank analysis was used to compare the results between the strains.

### The potential of the *aft4* locus as a transgene expression site for functional genomics applications

The results obtained so far suggest a significant potential of the *aft4* locus as a safe-haven site for functional genomics studies in *A. fumigatus*. To further explore this we examined heterologous expression of a nuclear targeted yellow fluorescent protein (NLS-Venus; (Nagai et al. 2002). The *venus*-encoding gene was placed under the control of a promoter which regulates the expression of the transcription factor *hapX* in an iron dependent manner (Schrettl et al. 2010; Gsaller et al. 2014) (Supplementary Fig. 4a). We generated two *phapX-venus* constructs, one with and one without the *hph* selection marker, Supplementary Fig. 4b) and introduced them into the *aft4* locus of the MFIG001 genome using CRISPR-Cas9 (Al Abdallah et al. 2017; van Rhijn et al. 2020). In agreement with our previous study where we targeted the *hph* selectable marker to the *aft4* locus (van Rhijn et al. 2020), all the tested transformants showed integration at the target site (8 out of 8 candidates) when we used marker-mediated transformation whereas 7% of growing colonies (1 out of 15 candidates) had a targeted insertion when we used the selection free approach (Supplementary Fig. 4c). These results demonstrate that the *aft4* locus can be used for efficient targeting of transgenes, especially in combination with our recently described selection-free CRISPR-Cas9 transformation system. We next characterized the phenotypes of the transformants in different culture conditions including iron-repletion and iron-starvation. Consistent with the results from the *aft4-hyg* knockout mutant (Fig. 2), no noticeable differences were observed in the growth profiles of the NLS-venus expressing mutants when compared to the isogenic control (Supplementary Fig. 5a and b).

Expression of the NLS-Venus protein was examined both at the protein and the RNA level. As shown in Fig. 4a, we were only able to detect nuclear localized yellow fluorescence from transformant strains when placed under iron-starvation conditions. Low iron induction of the NLS-Venus fusion protein and NLS-venus transcript was demonstrated by Western blotting analysis using a Venus specific antibody (Fig.4b) and qRT-PCR (Fig. 4c). We noticed that the Hyg-NLS-venus strain expressed a slightly higher level of venus mRNA (ca. 2-fold) compared to the NLS-venus stains perhaps as a result of the co-integration of the *hph* gene under the strong GDPA promoter (Supplementary Fig. 4b). The *aft4* locus is flanked by two putative ORFs (AFUB_092770 and AFUB_092780), which are arranged in a tandem orientation (Fig.1c) and in a relatively close proximity to *aft4*. We therefore investigated whether the insertion of the transgene expression cassette would have an impact on the expression of the neighbouring genes. As shown in Fig. 4d, the expression levels of the upstream ORF AFUB_092780 was not significantly impacted by insertion of the transgene however, the expression levels of the downstream ORF AFUB_092770 was expressed at a higher level in the NLS-venus strain, but not the hyg-NLS-venus strain, when compared to the isotype control. The reason for this change in expression of the neighboring gene was not investigated further however, no significant differences were observed in the growth characteristics of the NLS-Venus expressing strains in the corresponding culture conditions (Supplementary Fig. 5). Taken together, these results demonstrate the potential of the *aft4* locus for various functional genomics studies in *A. fumigatus*.

**Figure 4.**
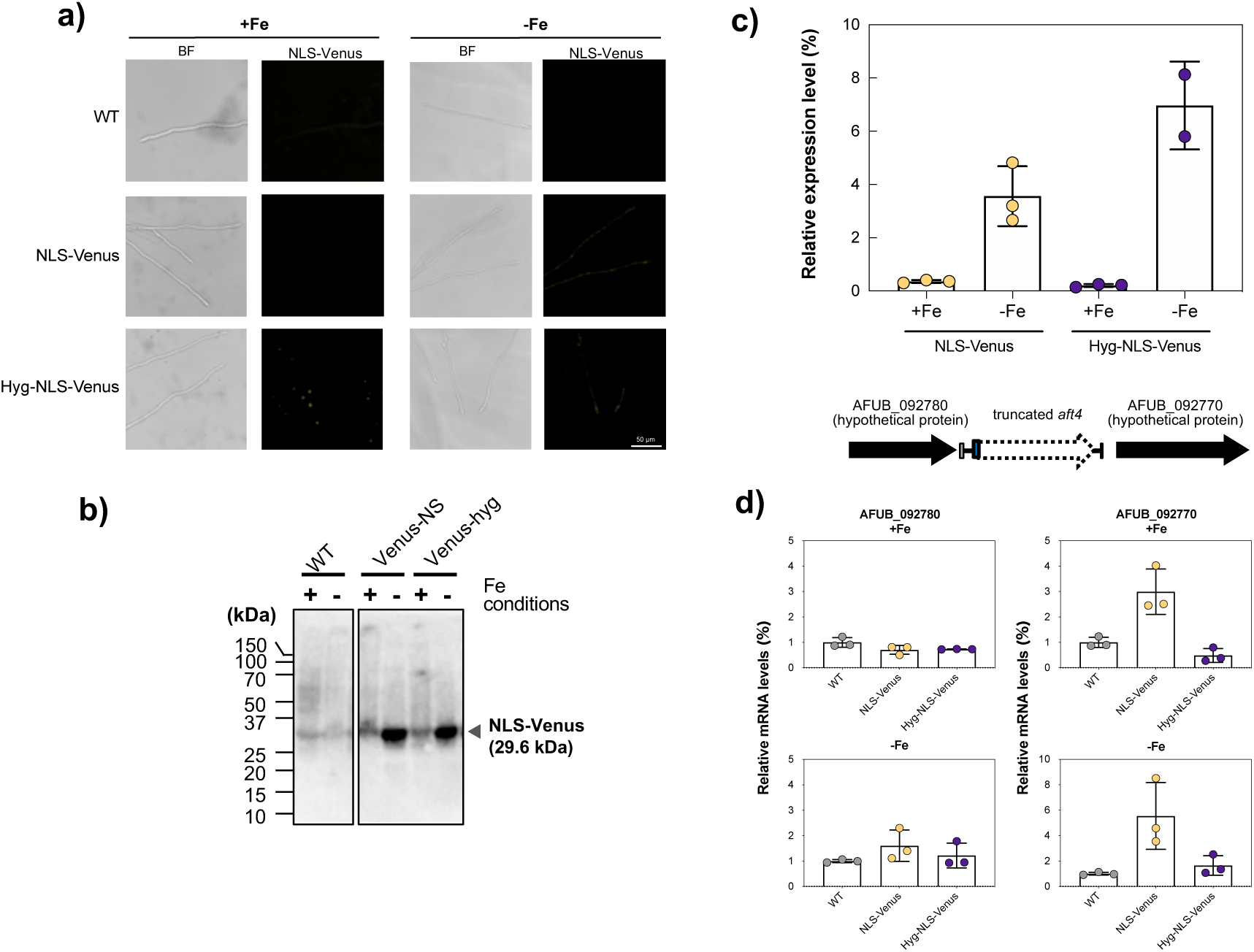
Heterologous expression of an NLS-Venus protein from the *aft4* locus. **(a)** Fluorescent microscopy of the NLS-venus expressing mutants (NLS-Venus and Hyg-NLS-Venus). The strains were grown in AMM for 24 h under iron-replete (0.03 µM; +Fe) or starvation (-Fe) conditions. **(b)** Western blot analysis showing iron level dependent inducible-expression of the NLS-Venus protein from the *hapX* promoter region. 100 µg of protein extracts were resolved on SDS-PAGE and probed for Venus protein using an anti-GFP polyclonal antibody. **(c)** Relative expression levels of *venus* transcripts in the NLS-Venus and the Hyg-NLS-Venus mutants after 18 h grown under iron-replete (0.03 µM; +Fe) or starvation (-Fe) conditions. The relative expression levels of *venus* transcripts were determined using the ΔΔCT method with the glyceraldehyde-3-phosphate dehydrogenase encoding *gdpA* as a reference gene. Data represent the mean of three independent biological experiments and error bars illustrates the standard deviation. Statistical significance was calculated by Student’s t test. **(d)** Transcript levels of the neighbouring genes (AFUB_092780 and AFUB_092770) in the wild-type (WT) and the NLS-Venus expressing mutants. Relative expression level of the neighbouring gene was determined as described above.

### The SH-*aft4* locus is highly conserved as a single-copy element in a large subset of clinical and environmental isolates of *A. fumigatus*

The fact that we couldn’t demonstrate the presence of the *aft4* locus in the genome of one of the commonly used laboratory strains of *A. fumigatus* D141 (Supplementary Fig. 2) prompted us to further investigate the conservation of the *aft4* locus in a large subset of *A. fumigatus* isolates. Using a recently published collection of sequence data (Sewell et al. 2019; Rhodes et al. 2021), we scrutinized the genomes of 218 different *A. fumigatus* strains that include 152 clinical and 63 environmental isolates. We identified 166 full length SH-*aft4* (the full-length safe haven including transposase and TIRs) homologs in 149 (68%) isolates. Through synteny analysis, we identified that *aft4* could be found in three distinct loci. Aft4 was positioned exclusively at locus NH1 in 132 strains, including Af293 and CEA10, at NH1 and NH2 in 16 strains and at NH1 and NH3 in 1 strain (Figure 5). The NH2 locus is found between two hypothetical genes that are absent in Af293 and CEA10 but share homology to the *Aspergillus lentulus* genes *IFM47457_08232, IFM47457_08230.* NH3 was flanked by one annotated gene, orthologous to AFUA_1G13970.

**Figure 5.**
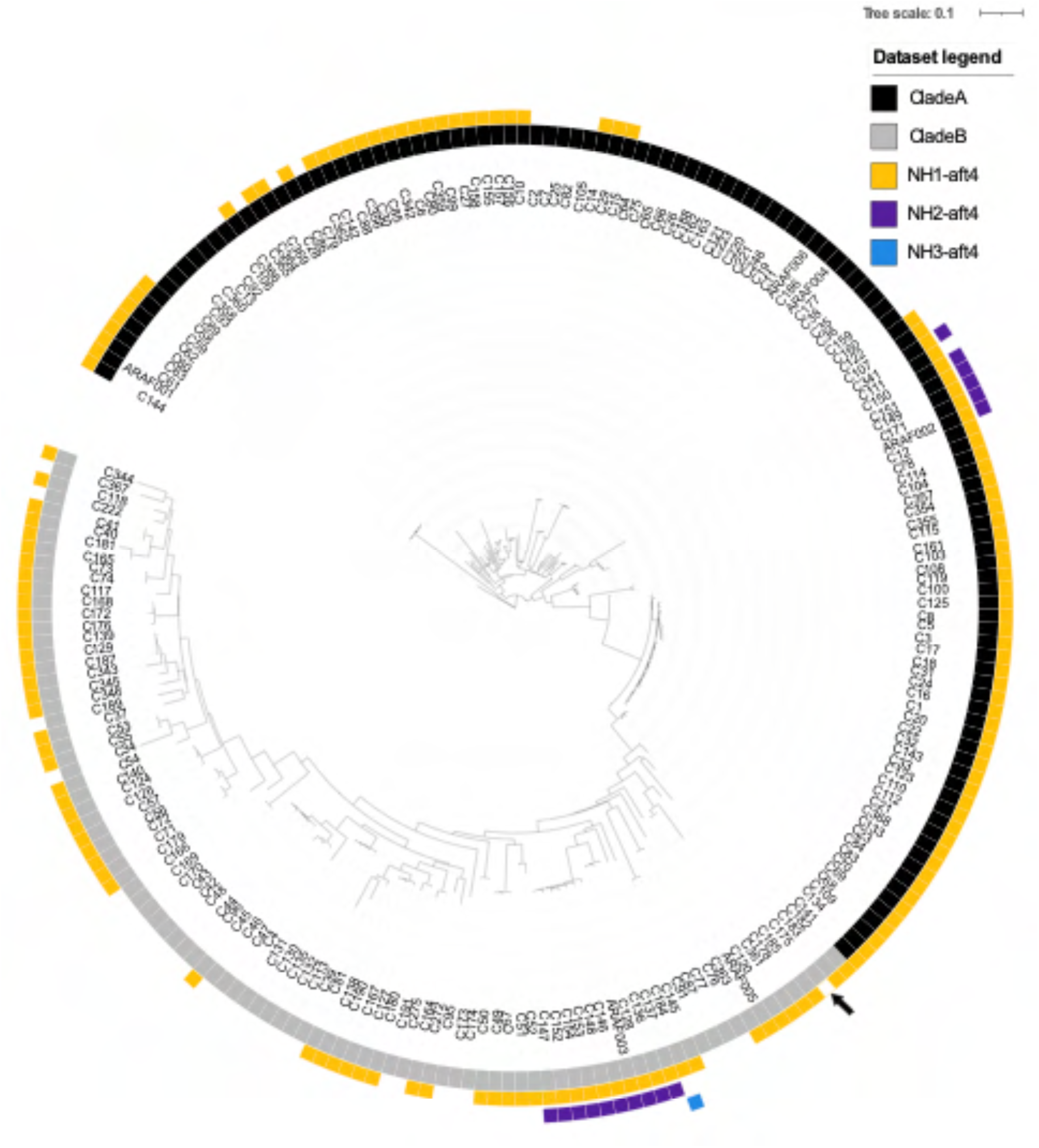
*aft4* is widely present as a single copy element in the genome of clinical and environmental isolates of *A. fumigatus*. Unrooted maximum likelihood phylogenetic tree (constructed in RAxML using genome-wide SNPs) showing the clade structure, A (black) and B (grey), and SH-aft4 loci/copy number, at regions NH1 (yellow), NH2 (purple) and NH3 (blue). Truncated AFUB_09770 at NH1 is highlighted in C175 (black arrow).

Nucleotide homology of *aft4* differed between genomic locations. At NH1*, aft4* had 98.9-99.5% sequence homology to the *aft4* transposase of Af293. In contrast, all *aft4* sequences found at NH2 were identical and had 91.3% identity to *aft4* of Af293 while the *aft4* orthologue at NH3 shared 94.3% identity. Extraction of potential protein encoding regions from these three loci show that all versions of *aft4* at NH1 and NH2 comprise truncated ORFs while *aft4* at the NH3*-*locus has the potential to produce a full-length, functional *aft4*-like transposase. Overall, these results indicate that the *aft4* locus is a highly conserved non-functional genetic region that exists as a single-copy element in the majority of *A. fumigatus* clinical and environmental isolates, and therefore the *aft4* locus could serve as a universal safe-haven site for most strains and genotypes of *A. fumigatus*.

## DISCUSSION

Identification of a well-defined genetic locus for the integration of a transgenic construct is an important step towards establishing a standardized approach for genetic complementation of a gene KO mutant as well as integration of a transgenic construct for various functional genomics studies. Genomic safe-haven sites (also known as genetic safe-harbor sites or the genetic safe-landing sites) have been identified in several organisms including a number of human fungal pathogens (Arras et al. 2015; Upadhya et al. 2017; Fan and Lin 2020; Pham et al. 2020). Recently two safe-haven regions (SH1 and SH2) were described in *A. fumigatus* located at intergenic regions of chromosome 1 and 2 of the Af293 genome (Pham et al. 2020) however no evidence was presented with respect to their potential impact on virulence in a mammalian system.

In this study, we explored the potential of an inactivated transposable element as a defined DNA integration site and identified the SH-*aft4* locus as a new genomic safe-haven site in *A. fumigatus*. Our genetic and biochemical analysis demonstrate that the deletion of the *aft4* element as well as the transgene expression from SH-*aft4* do not affect the growth characteristics and, in the case of *aft4* deletion, the pathogenic properties of *A. fumigatus*. We also provide evidence that SH-*aft4* can be used as a stable integration site for expression of a transgenic construct, and we have specifically shown that this site is compatible with selection-free CRISPR-Cas9 mediated transformation (van Rhijn et al. 2020). The *aft4* element was found to be highly conserved in the genomes of phylogenetically diverse clinical and environmental isolates indicating that SH-*aft4* locus can serve as an important molecular tool for genetic manipulation of *A. fumigatus* in a phylogenetically representative range of genotypes.

Our analysis suggests that the inactivated *aft4* element exist in the same genetic location in the genome of Af293 and A1163 and harbours the same nonsense mutations (Fig. 1b). Similarly, our diagnostic PCR revealed the presence of the *aft4* ORF and at least 200-bp of its flanking sequences in the other laboratory strain of *A. fumigatus*, ATCC46645 (Supplementary Fig. 2). As these strains are phylogenetically diverse, with 123 being in the recently describe clade A, and 95 in clade B, these facts might imply that the *aft4* transposable element was genetically inactivated and trapped at its current genetic locus at the early stage in the evolution of the *A. fumigatus* genome. Indeed, we observed that the *aft4* element is highly conserved as a single-copy element in a large subset of clinical and environmental isolates of *A. fumigatus* (Fig. 5). These results strongly suggest that the *aft4* locus can be used as a universal safe-haven site in not only the common laboratory strains but also in many other genotypes of *A. fumigatus*. Interestingly, however, we couldn’t confirm the presence of the *aft4* locus in the genome of the laboratory isolate D141 (Supplementary Fig. 2). Unlike the other common laboratory strains of *A.* fumigatus, D141 was derived from a patient with an aspergilloma (Staib et al. 1980), and several differences were observed in their biochemical and pathogenic characteristics compared to CEA10 and A1163 (Ries et al. 2019) indicating this strain may have undergone significant in-host adaptation leading to the possible loss of the SH-*aft4* site.

To summarise, we demonstrate the potential of the *aft4* locus as a transgene expression site for various functional genomics applications in *A. fumigatus*. We confirm that insertion of a 5.5 kbp of a gene expression construct can be introduced into the *aft4* locus with nearly 100 % efficiency in combination with a recently developed universal CRISPR-Cas9 methods (Al Abdallah et al. 2017; van Rhijn et al. 2020). We believe that the approach used in this study can provide a standardized strategy for genetic complementation of a gene knockout mutant as well as a novel locus to target expression cassettes or to perform studies that require a trans activation site.

## ACKNOWLEDGEMENT

This work was supported by the Medical Research Council (MRC) grant MR/M02010X/1 to MB and EB, the Biotechnology and Biological Sciences Research Council (BBSRC) grant number: 1640253 to MB and EB and the Wellcome trust grant 208396/Z/17/Z to MB. MCF and JR are supported by Wellcome trust grant 219551/Z/19/Z, NERC NE/P001165/1 and MRC MR/R015600/1

## COMPETING INTERESTS

MJB is a consultant to Synlab GmbH, is the director and shareholder of Syngenics Limited and is a substantive shareholder in PiQ Laboratories Ltd. The remaining authors declare no competing interests.

## SUPPLEMENTARY DATA

**Supplementary Table1:** Oligonucleotide used in this study.

**Supplementary Table2:** Characteristic features of the identified genetic safe-haven sites in *A. fumigatus*.

**Supplementary Table3:** Detailed characteristics of the SH-*aft4* in clinical and environmental *A. fumigatus* isolates

**Supplementary Figure 1:**
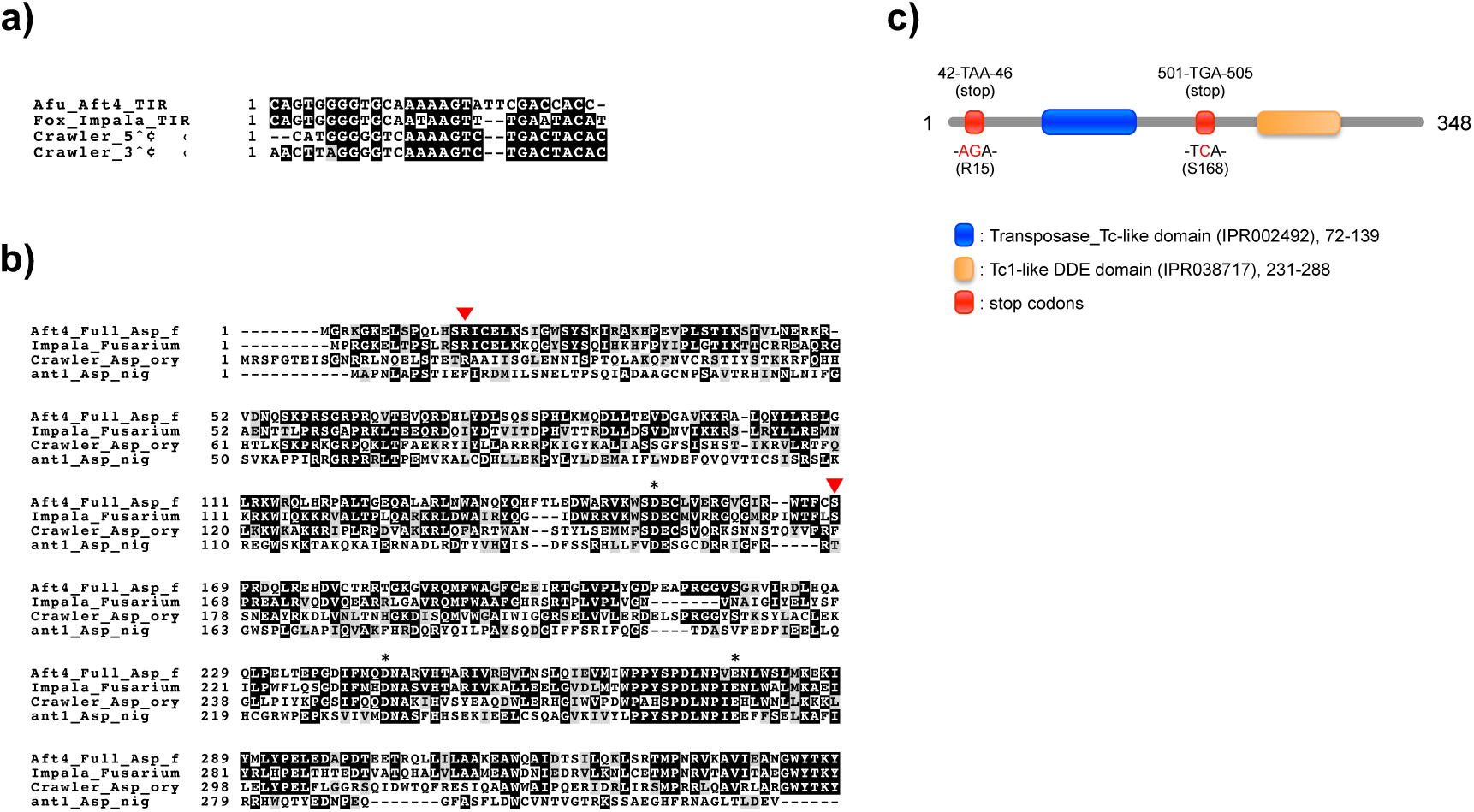
Domain structure and sequence alignment of the putative full-length form of Aft4 transposable element. **(a)** Alignment of Terminal inverted repeat (TIR) sequences of *A. fumigatus aft4*, *Fusarium oxysporum impala*, and *A. oryzae Crawler*. **(b)** Schematic representation of the domain structure of the putative full-length form of the *A. fumigatus* Aft4 transposase. The transposase Tc-like domain (IPR002492), and Tc1-like DDE catalytic domain (IPR038717) predicted by InterPro Scan is shown in blue and orange, respectively. The position of the stop codons found within the putative full-length ORF of *aft4* are shown in red with possible original codons. **(c)** Multiple protein sequence alignment of the transposase of the full-length form of the *A. fumigatus* Aft4, *F. oxysporum impala* (Langin et al. 1995), *A. oryzae Crawler* (Ogasawara et al. 2009)*, and A niger* Ant1 (Glayzer et al. 1995). The conserved amino acids consisting of the DD(35)E catalytic triad are indicated by asterisks. The position of the stop codons found in the inactivated *aft4* ORF are indicated by red triangles.

**Supplementary Figure 2:**
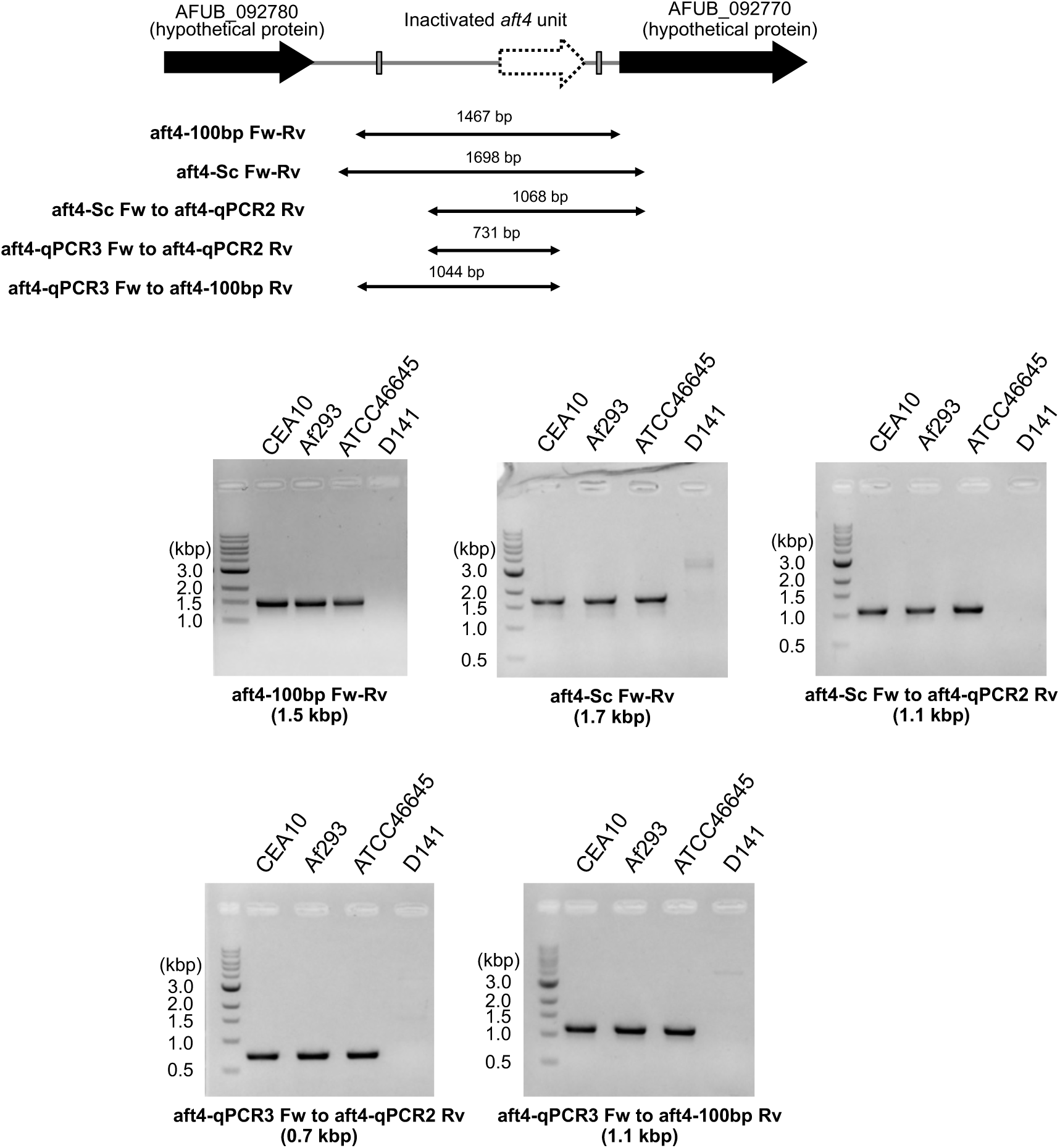
Diagnostic PCR analysis revealing the presence of the *aft4* locus in the genome of the *A. fumigatus* isolates in common laboratory use. **(a)** Primers used in the diagnostic PCR analysis are shown with the expected size of the PCR product, which is estimated from the *A. fumigatus* A1163 genome. **(b)** Results of the diagnostic PCR analysis. 10 ng of genomic DNA was used as a template for each PCR reaction and the generated PCR products were resolved on a 1% agarose gel. Positive amplification of the putative inactivated *aft4* ORF and the 1.7-kb region containing the ORF with 200-bp of the flanking sequences were observed for *A. fumigatus* Af293, CEA10 and ATCC46645.

**Supplementary Figure 3:**
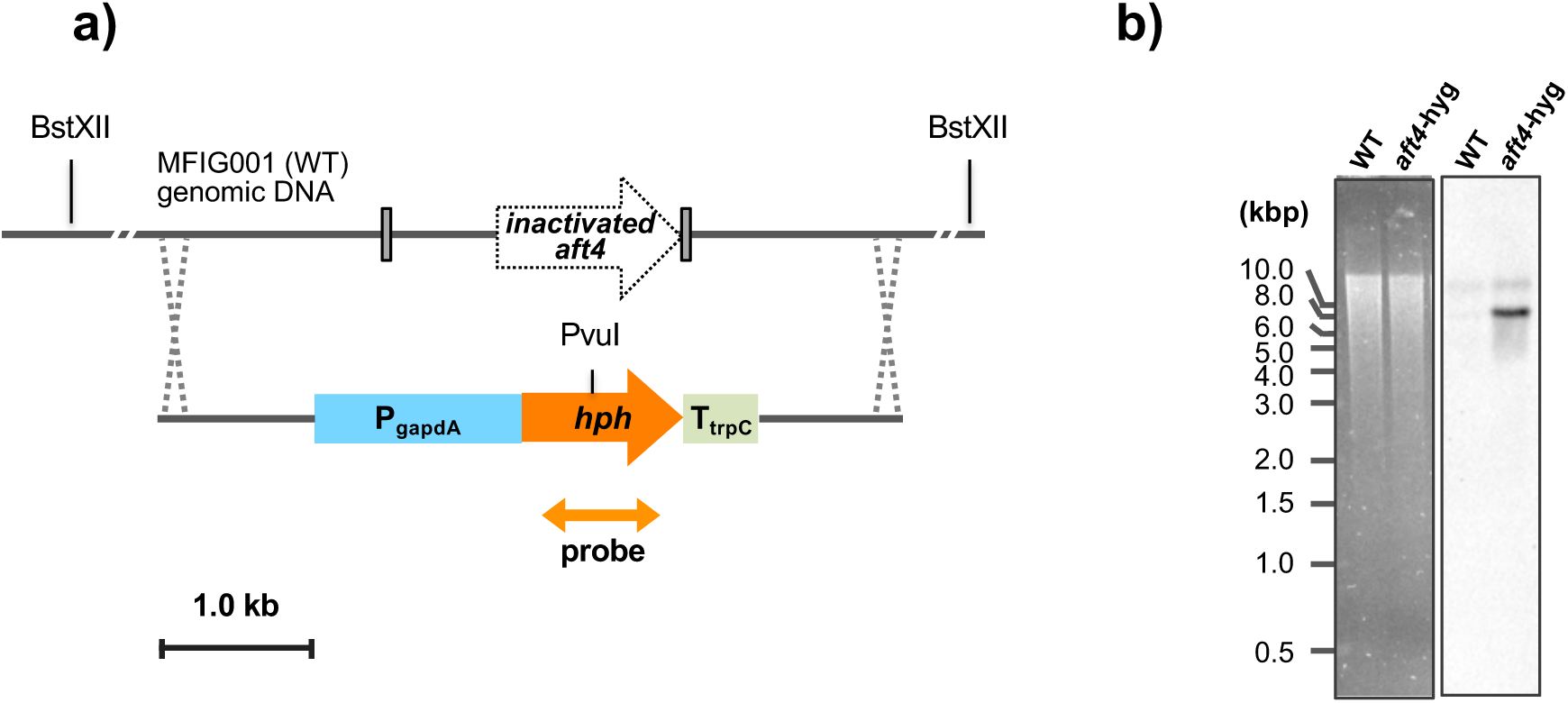
Southern blotting analysis verifying single copy integration of the hygromycine resistance (*hph*) cassette at the *aft4* locus. **(a)** Schematic representation of the construction of the *aft4* knockout mutant (*aft4-hyg*). The gene replacement cassette was constructed by fusion PCR and introduced into the wild-type *A. fumigatus* MFIG001 via homologous recombination. **(b)** Southern blot analysis of the *aft4-hyg* mutant. The restriction enzyme and the probe used in the Southern blot analysis are shown in the figure.

**Supplementary Figure 4:**
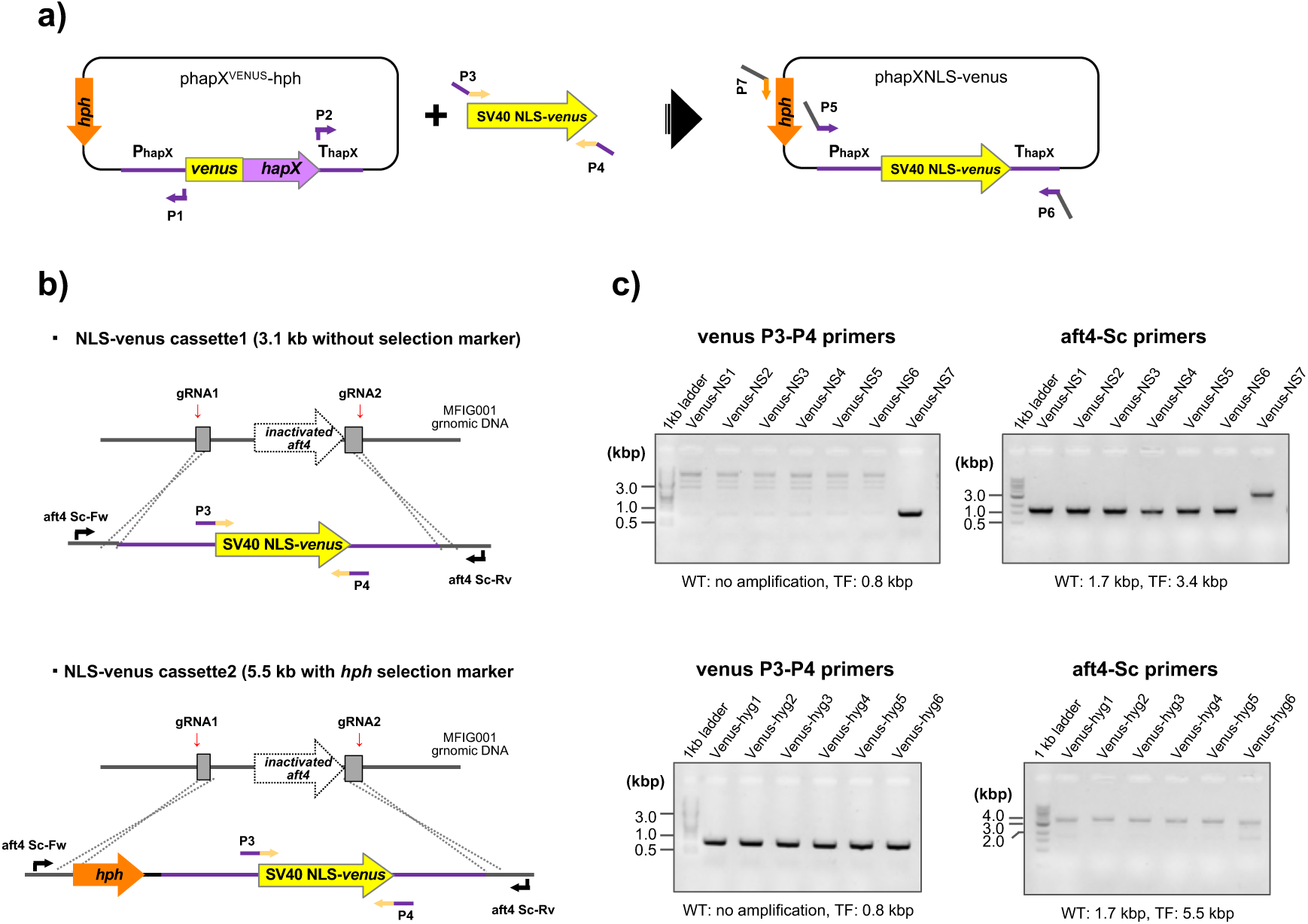
Construction of the NLS-Venus expressing mutants. **(a)** Scheme of the construction of NLS-venus expression cassettes for CRISPR-Cas9 mediated transformation. A DNA fragment containing 1.2 kb of the 5′- and 1 kb of the 3′-flanking regions of *hapX* derived from phapX^VENUS^-hph (Gsaller et al. 2014) was assembled with a SV40 NLS-venus fusion gene derived from pVenus-NLS (Furukawa et al. 2020) using DNA assembly. The selection-free expression cassette was amplified using the pair of primers P5 and P6, and the expression cassette with hygromycin resistance marker was amplified using the pair of primers P7 and P6, respectively. **(b)** Targeted integration of the NLS-venus expression constructs into the *aft4* locus using the CRISPR-Cas9 mediated transformation. Each NLS-venus expression cassette contains 50-bp of the homology arms to the corresponding guide RNAs (gRNA1 and gRNA2) to facilitate targeted integration at the *aft4* locus. The expression constructs were introduced to *A. fumigatus* MFIG001 by a CRISPR-Cas9 mediated genome-editing system. **(c)** PCR validation of the NLS-venus expressing mutants. Correct integration of the expression cassette was verified by PCR using the primer pairs venus-Sc Fw and Rv (for integration of *NLS-venus* fusion gene), and aft4-Sc Fw and Rv (for homologous integration). The primers used in the validation PCR are shown in **(b)** and Supplementary Table 1.

**Supplementary Figure 5:**
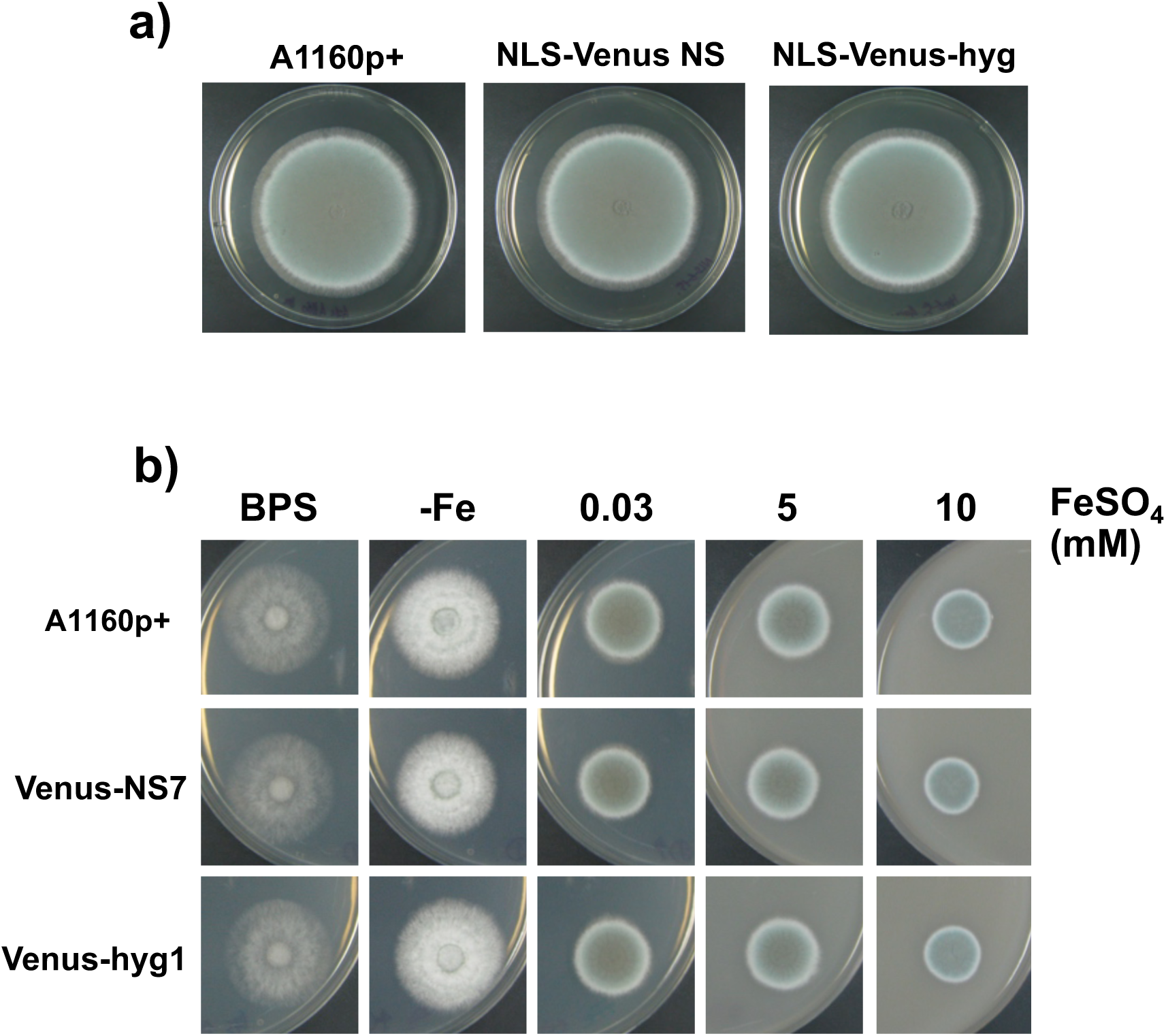
Growth characteristics of the NLS-Venus expressing strains on (a) ACM and (b) AMM with different iron conditions. 1×10^4^ spores were inoculated and the plates were incubated for 72 h at 37°C. -Fe media contains no iron, and BPS media contains 0.2 mM of the ferrous iron specific chelator bathophenanthroline disulfonate.

**Supplementary Figure 6.**
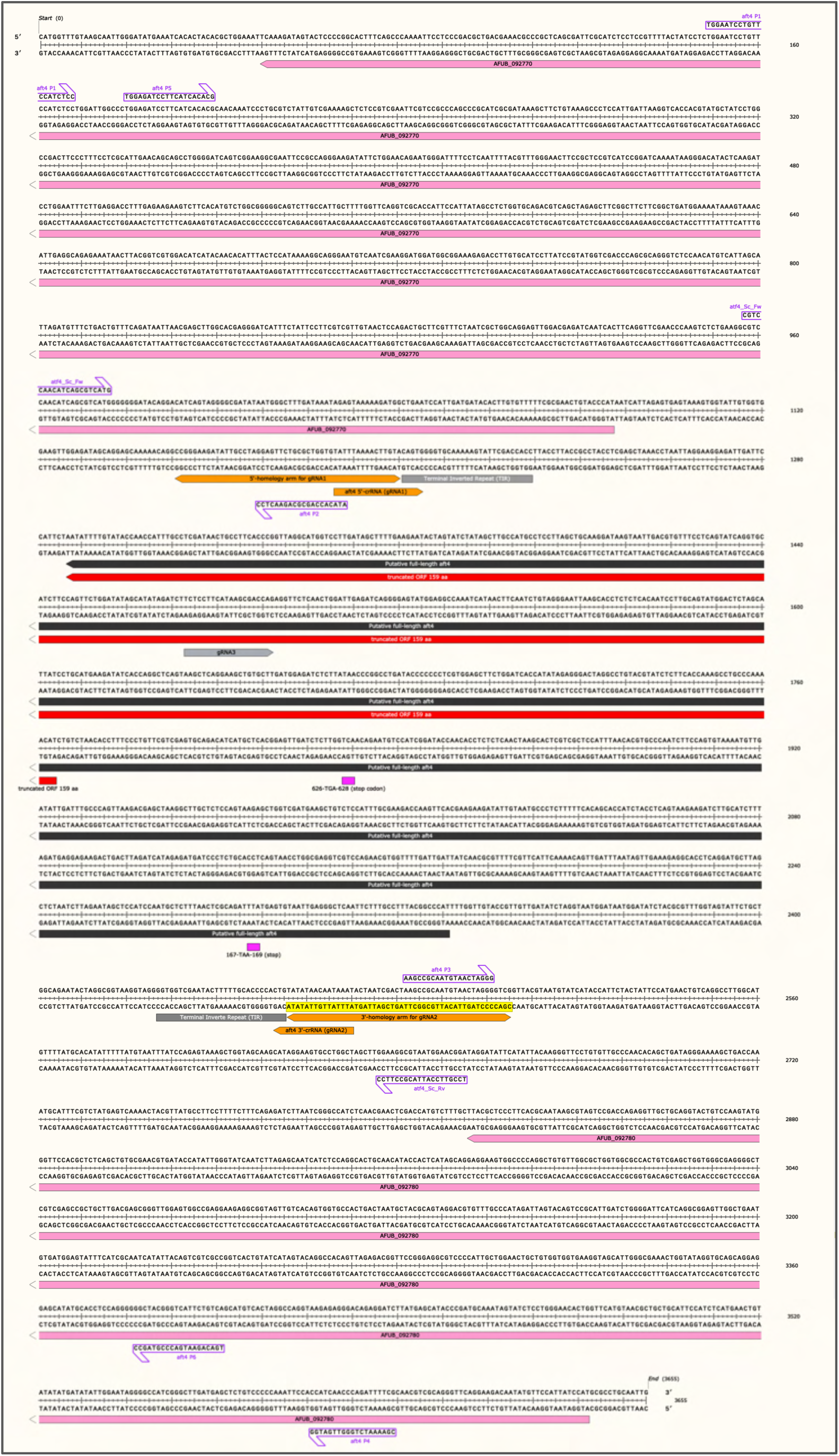
Primers and gRNAs used for genome editing at the *aft4* locus. The guide RNAs, the homology arms used to target the *aft4* locus, and the primers used to construct the NLS-venus expressing mutants are shown. Nucleotide sequence of the guide RNAs and the primers is also available in Supplementary Table 1.

## Notes

### Competing Interest Statement

I have read the journal's policy and the authors of this manuscript have the following competing interests: MJB is a consultant to Synlab GmbH, is the director and shareholder of Syngenics Limited and is a substantive shareholder in PiQ Laboratories Ltd. The remaining authors declare no competing interests.

